# Engineering a Juxtamembrane-Targeting CAR T-Cell Against Mesothelin: A Novel Binder Resilient to Shed Antigen in Ovarian and Pancreatic Cancer

**DOI:** 10.1101/2025.08.02.668276

**Authors:** John Scholler, Ai Song, Mansi Deshmukh, Decheng Song, Carl H June, Don L. Siegel

**Affiliations:** Center for Cellular Immunotherapies, Perelman School of Medicine, University of Pennsylvania, Philadelphia, USA; Department of Pathology and Laboratory Medicine, University of Pennsylvania, Philadelphia, USA; Parker Institute for Cancer Immunotherapy at University of Pennsylvania, University of Pennsylvania, Philadelphia, USA

## Abstract

Mesothelin is an attractive target for CAR-T therapy on a number of cancer types; however, the efficacy of this therapy is diminished because the bulk of the cell surface-expressed mesothelin is shed through naturally occurring proteolysis leaving behind a short juxtamembrane peptide “stump”. The two problems this creates are (1) the bulk of the target protein is no longer on the tumor cell and (2), the free soluble shed mesothelin remains in the tumor microenvironment and becomes present in blood/other body fluids binding to the mesothelin-targeted CAR-T and interfering with their ability to target the mesothelin that remains on the surface of the tumor. These issues have likely contributed at least in part to the lack of desired efficacy of CAR-T cells that target membrane distal regions of mesothelin (i.e., the shed domain) such as those utilizing anti-mesothelin scFvs SS1 and M5. Here we describe CAR T cells that utilize novel antibodies specific for the mesothelin stump domain thus being unaffected by the natural process of mesothelin shedding. Mesothelin “stump” specific CAR T cells (ST4-22) had cytotoxicity and *in vivo* activity that was comparable to standard anti-mesothelin CAR T cells. Importantly, CAR T cells expressing ST4-22 were effective against tumor cells that were resistant to standard anti-mesothelin CAR T cells.

## Introduction

Mesothelin (MSLN) is a glycosylphosphatidylinositol (GPI)-anchored cell surface protein usually restricted to mesothelial cells but is overexpressed in multiple solid tumors, including ovarian and pancreatic cancers (1–3). Its tumor-specific expression and limited presence in normal tissues make it an attractive target for antibody and chimeric antigen receptor (CAR) T-cell therapy (4–13).

However, development of effective anti-mesothelin CAR T cells has been challenged by dense stroma (14, 15) and the presence of shed mesothelin in the tumor microenvironment (16). Soluble mesothelin can act as a decoy, binding CARs and hindering T-cell engagement with tumor cells (16–18). To overcome this, Liu et al. engineered CAR T cells targeting a juxtamembrane epitope near the membrane protease cleavage site of mesothelin using the 15B6 antibody (19). These CAR T cells retained activity in the presence of shed antigen and exhibited potent antitumor efficacy in preclinical models. In a subsequent study, a humanized version (h15B6) demonstrated marked activity against pancreatic cancer xenografts, overcoming resistance observed with shed-binding CARs (20).

Beyond mesothelin, alternative tumor-associated antigens are being explored. CEACAM7, expressed in pancreatic ductal adenocarcinoma (PDAC) but absent from most normal tissues, was recently validated as a promising CAR T-cell target. Raj et al. developed CEACAM7-directed CAR T cells that effectively eliminated CEACAM7-positive PDAC cells in vitro and induced remission in patient-derived xenograft models without observable off-tumor toxicity (21).

Building on these advances, we here describe a novel CAR T-cell construct targeting a previously unexploited epitope within the juxtamembrane region of mesothelin. Our approach leverages insights from prior studies to avoid inhibition by shed mesothelin, while improving tumor specificity and functional activity in ovarian and pancreatic cancer models. We detail the design, functional characterization, and antitumor efficacy of this new CAR T-cell, highlighting its potential to overcome current therapeutic limitations.

## Results

### Isolation and characterization of scFvs targeting membrane proximal epitopes of mesothelin

Membrane-bound mesothelin (MSLN) protein (**Fig. 1**) is derived from a 622-amino acid precursor protein which subsequently undergoes a series of processing steps that results in a 303-amino acid mature form of the protein (amino acids 296-598) attached to the cell surface through a GPI linkage (19, 22, 23). Mesothelin can be shed from the cell membrane through the action of proteolytic enzymes with major cleavage sites within a carboxy terminal peptide comprising amino acids 582 to 598 (19, 20, 24). This results in the release of the bulk of mesothelin beginning at residue 296 and extending to approximately residue 581, and retention of a membrane-bound mesothelin “stump” domain beginning around residue 582 and extending to residue 598 (**Fig. 2A**).

**Fig. 1.**
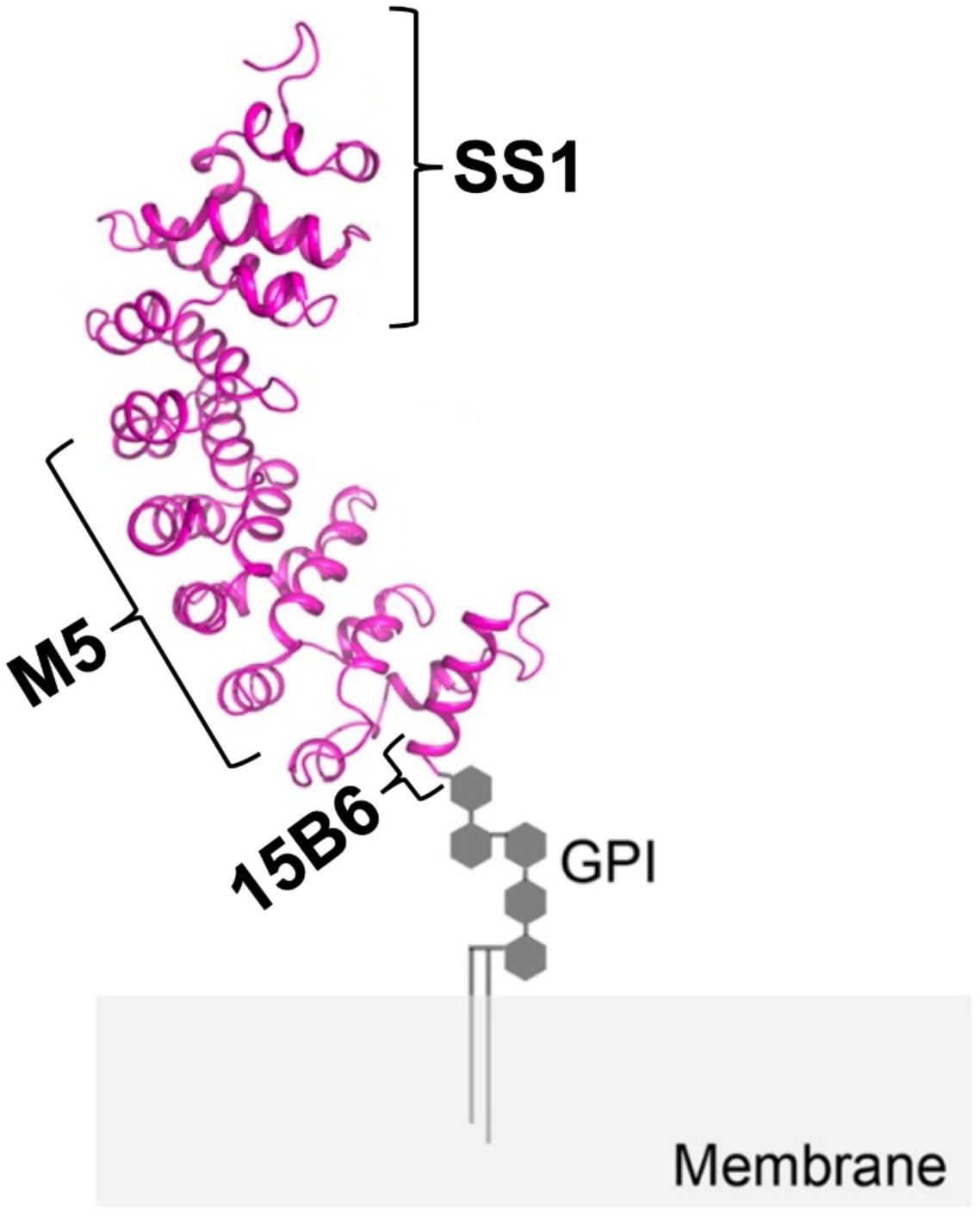
Schematic of mesothelin. Cartoon showing mesothelin and its GPI linkage, and binding sites for 15B6, M5 and SS1 antibodies. M5 and SS1 bind at sites distant from the MSLN proteolytic cleavage sites and can be released by shedding and can also bind to shed MSLN. The 15B6 scFv binds to cell-associated MSLN stump near the plasma membrane which is not released.

**Fig. 2.**
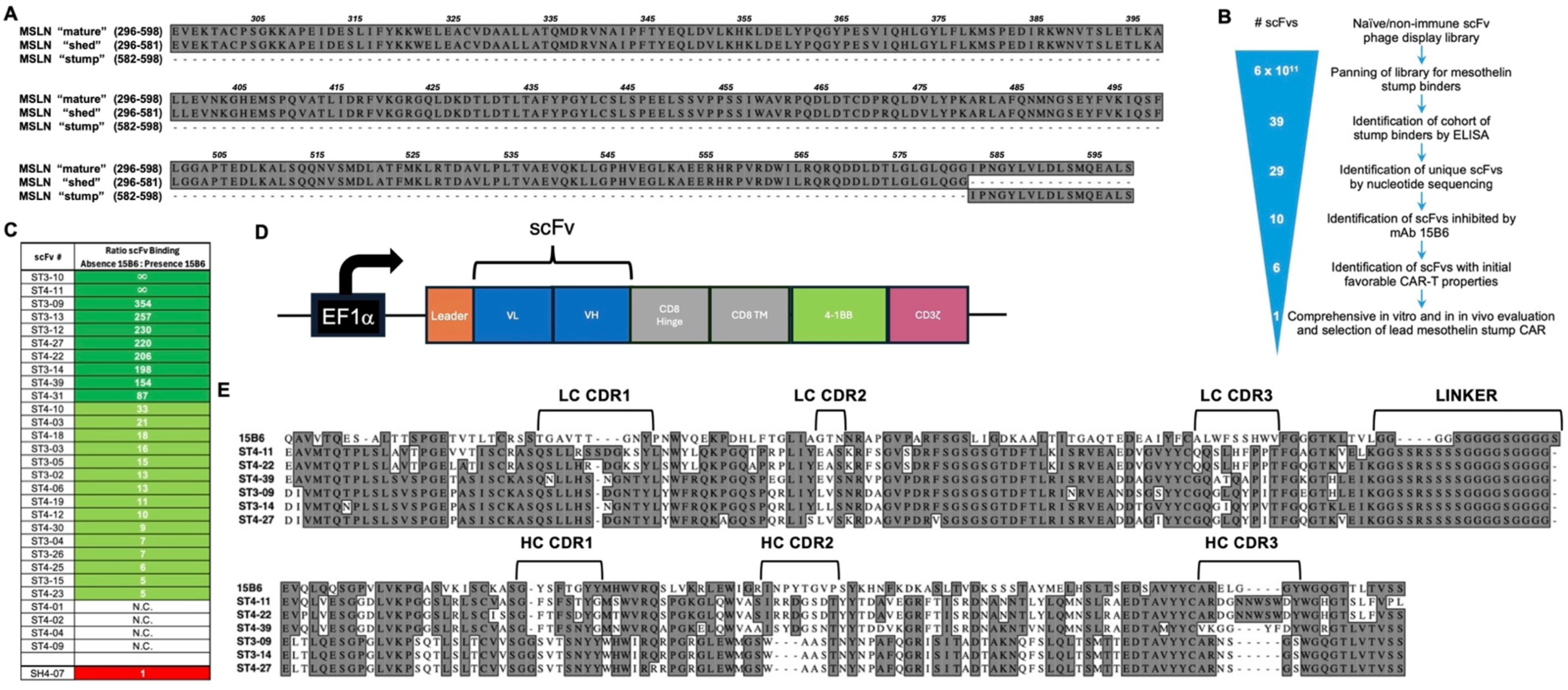
Isolation and characterization of scFvs targeting membrane proximal epitopes of mesothelin. (**A**) Amino acid alignment of mature, shed, and stump region MSLN proteins. (**B**) Canine B-cell derived scFv naïve/non-immune phage display library screening strategy. Positive/negative panning on full-length MSLN and shed-length MSLN proteins, respectively, enriched for scFvs targeting stump region of MSLN. Twenty-nine unique stump binders identified by nucleic acid sequencing from an initial cohort of 39 were further characterized as scFvs and a subset as T cell chimeric antigen receptors. (**C**) Competition ELISA assay used to rank ability of mAb15B6 reference MSLN stump-directed scFv to block binding of each of 29 stump-directed canine scFvs. Ratios of canine scFv binding to MSLN in the absence and presence of 15B6 are tabulated. The top 10 blocked scFvs (dark green shading) were chosen to be further screened as chimeric antigen receptors in order to narrow down the number of candidates for comprehensive evaluation to a total of 6. For 4 of the 29 scFvs, ELISA signals were too low to confidently determine binding ratios (“N.C.”). ScFv SH4-07, a scFv from library panning determined to be specific for the shed region of MSLN, served as a negative control for assay displaying equal binding to full-length MSLN whether 15B6 was present or not (ratio = 1). (**D**) Schematic of CAR construct inserted downstream of an EF1α-derived promoter within a 3rd-generation lentiviral vector plasmid comprising a CD28 transmembrane domain and 4-1BB and CD3ζ signaling domains. (**E**) Amino acid alignment of 15B6 reference MSLN stump-directed scFv with 6 canine stump-directed scFvs evaluated CARs in this study. All scFvs were used in variable light chain/variable heavy chain (VH) orientation. 15B6 scFv sequence and linker per Liu et al. (19, 24). Linker for canine scFvs as per pComb3X phage display vector system (25). Light chain and heavy chain CDR assignments per IMTG nomenclature and identified using Domain-GapAlign tool (44).

Antibody binding fragments specific to the stump region of MSLN were isolated from a canine scFv naïve/non-immune IgM/IgG/λ/κ phage display library containing an estimated 40 billion individual transformants constructed from canine B cell mRNA and the pComb3X phagemid vector as described (25–27) using oligonucleotide primers based on published canine immunoglobulin heavy and light chain germline genes (28).

In our experience, canine immunoglobulin genes show relatively high homology to human counterparts, and scFvs isolated from this library have been useful in not only veterinary clinical applications but in a human clinical trial (www.clinicaltrials.gov NCT06544265).

To isolate scFvs to the stump region of MSLN, the scFv phage display library was panned using solid phase technique against immobilized full-length recombinant MSLN protein after negative selection against a recombinant form of MSLN representing shed MSLN. Negative selection to shed domains was achieved through two means: (1), pre-incubation of the phage library in streptavidin-coated microplate wells with immobilized biotinylated MSLN (296–580) and subsequent retrieval of non-absorbed phage particles and (2), addition of soluble non-biotinylated MSLN (296–580) to the non-absorbed phage particles to serve as a competitor during positive selection in wells coated with streptavidin and full-length (296–598) recombinant biotinylated MSLN.

Starting with ∼600 billion scFv phage clones, after 4 rounds of negative/positive selection, ∼400 million potential stump binders were isolated from panning round outputs. Sixty-six scFv phage clones were randomly selected from panning rounds 3 and 4 (clone nomenclature comprising “PX-Y” where “X” is panning round and Y is arbitrary clone number), produced as monoclonal phage-displayed scFvs, and examined by phage ELISA for binding to the truncated and full-length forms of MSLN (**Fig. S1**).

Of the 66 randomly selected scFv phage clones, 39 bound to full-length MSLN but not truncated MSLN suggesting specificity for the stump domain which is present in the long form but not the short form (**Fig. 2B**). Of these 39 presumed “stump-binders”, nucleotide sequencing of the scFv heavy and light chain variable regions revealed 29 unique scFvs. Of the remaining 27 randomly selected scFv phage clones, 26 bound to both long and truncated MSLN suggesting specificity to the shed region of MSLN, and 1 clone bound to neither. Of these 26, 7 of them were unique scFvs based on nucleotide sequencing of heavy and light chains. Having binned scFvs to MSLN stump binders vs. shed MSLN binders, scFv clone names were modified by removing “P” and replacing with “ST” or “SH”, respectively, to represent “STump” or “SHed”.

To reduce the number of unique stump binding scFvs to be evaluated *in vitro* and *in vivo* as T cell chimeric antigen receptors, two sequential screenings were performed. First, the extent to which reference murine mAb 15B6 could inhibit the binding of each of the 29 scFvs to full-length mesothelin was determined to assess stump epitope specificity.

To develop this assay, a recombinant form of 15B6 was produced as a full-length murine IgG1 using the published amino acid sequences for the 15B6 heavy and light chains (19).

After verifying that recombinant 15B6 retained its binding to the mesothelin stump domain (**Fig. S2**), microplate wells coated with full-length MSLN were preincubated (or not) with an amount of 15B6 enough to saturate the MSLN coating the wells. Phage displayed scFvs were then applied to MSLN-coated wells (with or without bound 15B6) and after washing away unbound phage, bound phage were detected using an HRP-labeled anti-M13 phage secondary reagent (**Fig. S3**). For each scFv, the ratio of scFv binding in the absence and presence of 15B6 was calculated **(Fig. 2C)** and the 10 (of 29) scFvs most strongly blocked by 15B6 were selected for further study in a second screening assay. A useful internal control was scFv P4-07 (which was determined to bind to shed MSLN, **Fig. S1**, and renamed SH4-07, **Fig. 2C**) which showed a 15B6 inhibition ratio of unity as would be expected from a MSLN shed-directed scFv.

For the second screening assay, the 10 scFvs selected from the 15B6 inhibition assay were cloned into our CAR construct (**Fig. 2D**), and human T cells or Jurkat NFAT reporter cells were transduced. A limited series of functional tests to assess the degree of tonic signaling, peak activation, and *in vitro* killing of mesothelin-expressing cell lines was performed (data not shown). The results of this additional scFv screening assay enabled us to reduce our cohort of MSLN stump-directed scFvs from 10 candidates to 6 candidates which displayed acceptable performance properties (**Fig. 2B**).

Alignment of the predicted amino acid sequences for the final group of 6 scFvs along with 15B6 reference scFv (**Fig. 2E**) show that all 6 scFvs are quite distinct from 15B6. All utilize kappa isotype light chains. ST4-11 and ST4-22 show identical CDR2 and CDR3 regions in both heavy and light chains but have marked differences in their CDR1 regions as well as framework regions 1 and 4. ST3-09, ST3-14, and ST4-27 have nearly identical heavy chains (except for a single amino acid difference in framework 2 of ST4-27), but they are paired with light chains that show significant degrees of diversity in all framework and CDR regions. ST4-39 is distinctly different in primary structure from the other 5 scFvs (**Table S1)**.

### Screening top six stump CARs

The top six stump CAR scFv candidates were engineered into healthy (normal donor, ND) human T cells using lentiviral vector technology for *in vitro* and *in vivo* evaluations to determine their ability to target specifically and control tumor growth of cells expressing MSLN as either the full length or just the stump portion and confirming that recognition was MSLN specific. Three different tumor cell lines were evaluated: AsPC1 (pancreatic ductal adenocarcinoma), H838 (non-small cell lung adenocarcinoma) and OVCAR8 (ovarian adenocarcinoma). The percentage transduction of each stump CAR in ND608 donor T cells and their levels of expression (MFI) can be seen in **Fig. S4**. The performance of each stump CAR was compared relative to our conventional MSLN targeting M5 CAR and the non-MSLN targeting PSMA or anti-CD19 hCART19/CTL119 CARs. The M5, PSMA and hCART19/CTL119 CARs have been previously described (29–31).

To determine relative efficacy and target specificity, each stump CAR was co-cultured with GFP expressing H838 or AsPC1 cell lines expressing either their normal full length MSLN (wild type, WT), only the stump form of MSLN, or no MSLN (MSLN-KO) for 7 days (**Fig. 3A, B or C**, respectively). Killing efficiency was measured by Incucyte® live cell imaging every 3 h for changes in GFP intensity, which directly correlate to number of tumor cells surviving. All stump CAR T cells co-cultured with wild type AsPC1 cells impaired the tumor growth with only ST4-11 and the non-targeting PSMA CAR unable to show anti-tumor efficacy. Stump CAR T cells co-cultured with the wild type H838 cells were able to control tumor but with a wider range of effectiveness. ST4-22, 15B6, ST4-39 and M5 had the ability to reduce the tumor numbers, while ST3-09, ST3-14, ST4-11, ST4-27 were only able to maintain steady tumor cell numbers in this faster growing cell line (**Fig. 3A**). These results indicated that the stump CARs could control full length MSLN comparably to our standard CAR M5 that binds to an epitope distal to the stump epitope (29).

**Fig. 3.**
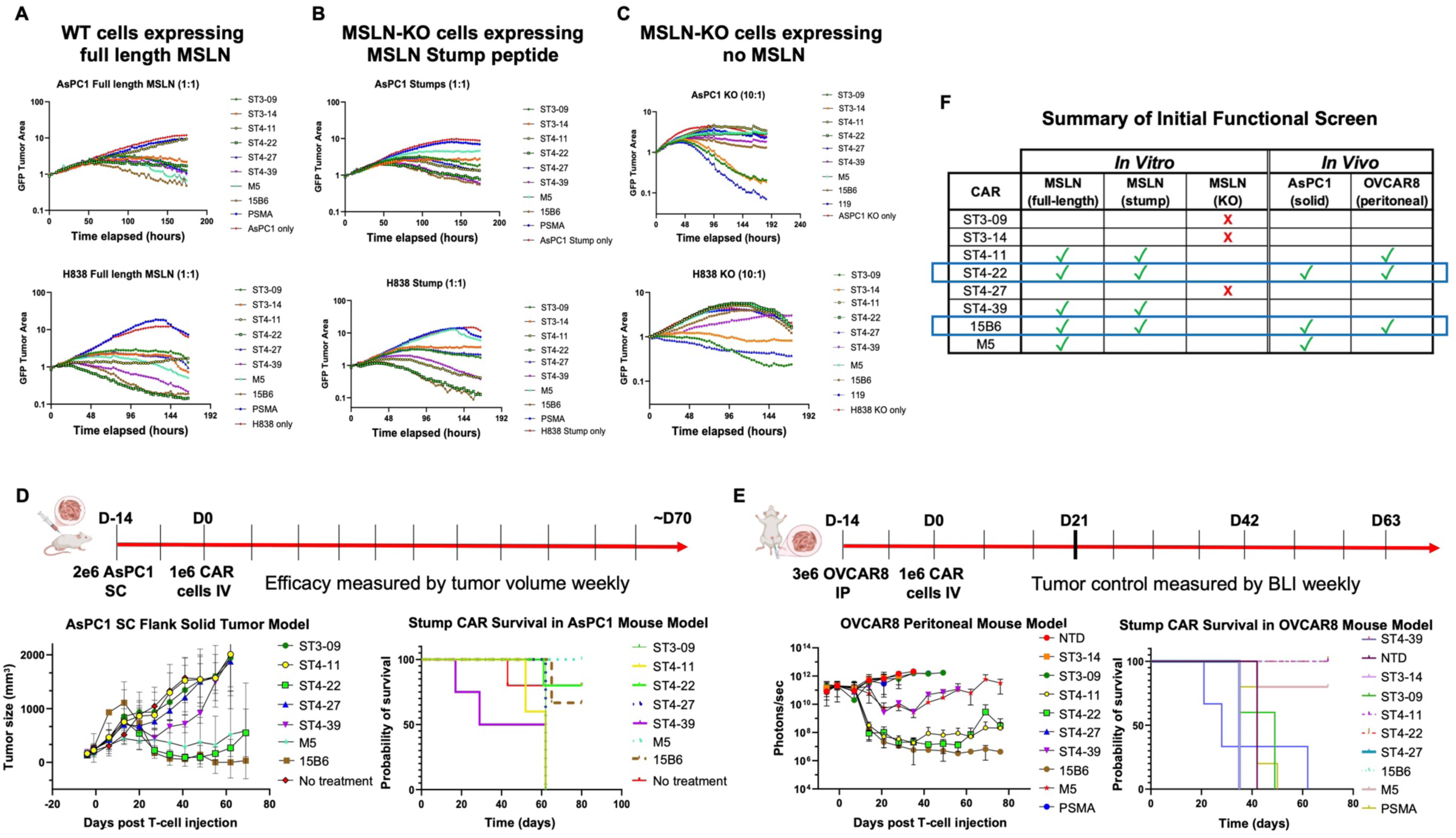
Stump CAR targeting and efficacy screen in healthy donor ND608 T cells. (**A**) AsPC1 and H838 wild-type cells expressing full length MSLN were co-cultured with stump CAR T cells at a 1:1 E:T ratio and cytotoxicity measured by Incucyte®. (**B**) Stump CAR T cells were co-cultured with AsPC1 and H838 cells at a 1:1 E:T ratio where the MSLN gene was knocked out (KO) and transduced with lentiviral vector to only express the MSLN stump peptide. (**C**) AsPC1 and H838 MSLN-KO cells were co-cultured with the stump CAR T cells at a 10:1 E:T ratio to assess for non-specific target recognition. (**D**) Schema and results for *in vivo* screen of stump CAR tumor clearance in AsPC1 NSG mouse models. 14 days after implanting 2e6 AsPC1 cells in right flank, 1e6 CAR T cells were injected IV. Tumor sizes were measured weekly and survival recorded (5 mice per group). (**E**) Schema and results for *in vivo* screen of stump CAR tumor clearance in OVCAR8 NSG mouse models. 14 days after intraperitoneal injection of 3e6 OVCAR8 tumor cells, 1e6 CAR T cells were injected IV by tail vein (5 mice per group). Weekly tumor BLI progression and survival was recorded. (**F**) Summary of initial *in vitro* and *in vivo* functional screens of MSLN stump-direct CARs.

To confirm the specificity of the stump CARs to the stump region of MSLN GPI linked surface expressed protein, we first knocked out the MSLN expression (MSLN-KO) in both the AsPC1 and H838 cell lines by CRISPR technology. These MSLN-KO cells were then used to identify stump CAR T cells that could kill tumor cells independent of stump targeting. In addition, to confirm that stump CAR T cell killing was directed specifically to the stump portion of MSLN, we transduced these MSLN-KO cell lines with a lentiviral construct encoding the 582-598 amino acid membrane proximal stump region (**Fig. 2A**), which included a 3x FLAG tag on the amino terminal end for detection. Stump CARs co-cultured with AsPC1 cells that express the stump epitope showed that each of the stump CARs slowed tumor growth with ST4-22, ST4-39 and 15B6 showing the greatest reduction (**Fig. 3B, top**). As expected, CAR M5 lost its ability to control tumor cells only expressing the stump epitope. Similarly, in H838 co-cultures, ST4-22 and 15B6 recognized the stump region the best, followed by ST4-11 and ST4-39 (**Fig. 3B, bottom**).

Next, we co-cultured the anti-stump CAR T cells with AsPC1 or H838 MSLN-KO cells at a higher 10:1 E:T ratio to identify non-specific MSLN CAR T cell activity. CAR T cells expressing the ST3-09, ST3-14 and ST4-27 scFv had non-specific killing activity on both cell lines (**Fig. 3C**).

In separate experiments, we transduced our stump CARs into Jurkat NFAT-GFP reporter cell lines to evaluate ligand-independent signaling (**Fig. S4D**). The induction of NFAT signaling was evaluated on wild type or MSLN-KO H838 cells. Stump ST4-22 and 15B6 CAR T cells had ligand dependent signaling, which was specific to MSLN expressing cells, similar to our distal targeting M5 CAR T cells expressed in the Jurkat reporter cell line.

The stump CAR activities were next evaluated *in vivo* using the xeno-mouse models of the pancreatic AsPC1 flank solid tumor and the ovarian OvCAR8 peritoneal tumor. In the AsPC1 model after 2 weeks of engraftment, 1e6 stump CAR T cells were administered IV and changes in tumor growth were measured weekly (**Fig. 3D**). By three weeks, ST4-22, 15B6 and M5 CAR T cell treated mice had potent antitumor effects and most mice were able to eliminate visual detection of the tumor with long-term survival, with a few animals succumbing to xenogeneic GVHD. ST4-39 showed early antitumor effects but failed to have persistent control tumor growth by 4 weeks.

The tumor growth kinetics in individual mice treated with ST4-11, ST4-22, 15B6 and M5 CAR T cells are shown in **Fig. S4B**. All other stump CARs failed to show any signs of tumor reduction. Based on these results, ST4-22 was the leading candidate in the AsPC1 pancreatic tumor model.

The OvCAR8 tumor cell line is known to produce proteases which cleave large amounts of cell surface MSLN (19, 24), producing a milieu of shed soluble antigen capable of blocking MSLN CAR T cells, reducing their ability to target surface antigen, and kill tumor. Thus, the OvCAR8 peritoneal mouse model is suitable for evaluating the ability of the stump CARs to bypass resistance caused by cleaved MSLN to target the tumor cells, engaging with the surface-bound full length MSLN as well as the residual membrane proximal MSLN stump remaining after proteolytic cleavage.

Two weeks after peritoneal implantation of OvCAR8 tumor cells, 1e6 stump CAR T cells were IV administered and by two weeks only the stump CARs ST4-22, ST4-11 and 15B6 had potent antitumor efficacy, leading to long-term survival (**Fig. 3E**). It is notable that CAR M5 was able to effectively control AsPC1 tumors in our previous experiments. **Fig. 3F** summarizes the *in vitro* and *in vivo* screening of the top six stump CAR candidates. These results indicate ST4-22 as our top candidate having good efficacy in both tumor models, including the resistant OvCAR8 tumor, and with no background ligand-independent signaling activity and no non-specific killing of MSLN KO cells.

To further evaluate the stump CAR T cells, we evaluated ST4-22 with three separate normal T cell donors covering both sexes and across ages from 27 to 62 years. Before this stringent evaluation, we first minimized potential for induced immune responses to our CAR T cells, by eliminating internal ATG start sites from alternate reading frames encoded in the transgene that would produce translated peptides greater than 50 amino acids in length. By mutating ATG start sites and not changing the amino acid composition of the CAR, we generated a modified nucleotide sequence of ST4-22 which we now designate CAR 422 (**Fig. S5 and Table S2**)

The manufacturing profiles of CAR 422 in the healthy donors ND630 (27 yo female), ND410 (61 yo female), and ND616 (62 yo male) are similar in doubling rates and size compared to 15B6 and control CARs M5 and CAR 119 (**Fig. S6**). The killing efficiency of the CAR T cells from each of the three donors was first evaluated in *in vitro* co-cultures with AsPC1 and OvCAR8 tumor cells at CAR T cell to tumor cells of 10:1, 3:1, 1:1 and 0.3:1. Targeting the AsPC1 cells, CAR 422 T cells had killing efficiency of greater than 75% except for the lowest E:T ratio of 1:3 in donor ND630. Overall, the efficacy of tumor control was slightly less than the stump CAR 15B6 and the distal targeting CAR M5 (**Fig. 4A**). In the co-cultures with OvCAR8 tumor cells, CAR M5 maintained a stable tumor control down to a 1:1 ratio, while both stump CARs 422 and 15B6 lost tumor control at the 1:1 ratio for ND630 and ND616 and at the 1:3 ratio for ND410 (**Fig. 4B**). In all co-cultures, CAR 422 showed a slightly lower efficacy profile compared to 15B6. The robustness of the CAR M5 in *in vitro* OvCAR8 co-cultures compared to the *in vivo* results shown in Figure 3 probably reflects the time needed to build up the effective blocking concentrations of shed material in the tumor microenvironment. In the co-cultures, the shed material is likely just beginning to build up while in the OvCAR8 IP model, the shed material has already been accumulating for several weeks and may be more representative of a patient’s tumor microenvironment.

**Fig. 4.**
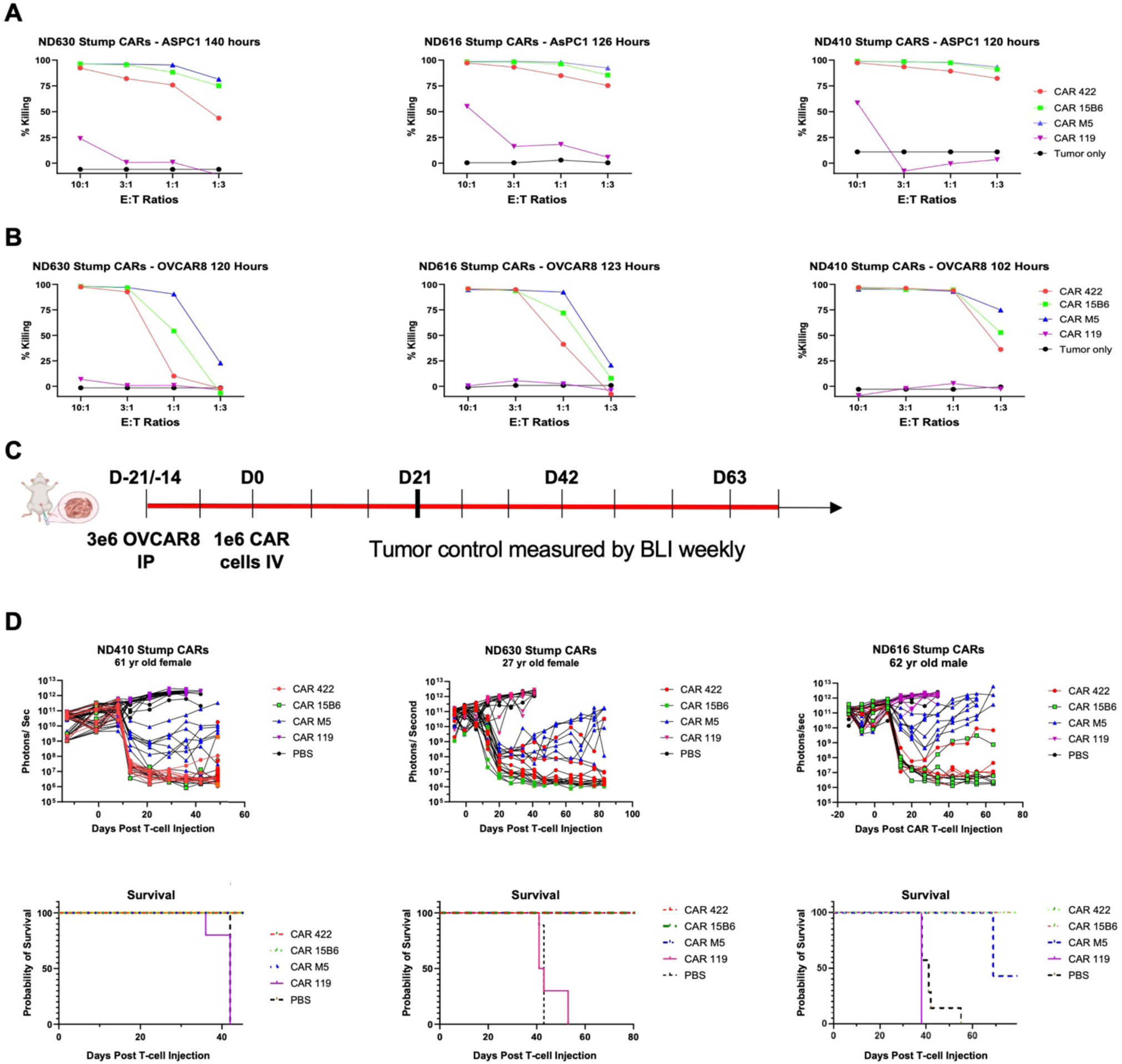
Killing efficacy of stump CARs *in vitro* and *in vitro* across multiple T cell donors. T cells from 3 healthy donors varying in gender and age were transduced with MSLN stump-directed CAR 422 and CAR 15B6, along with distal MSLN targeting CAR M5 and CD19-targeting CAR 119 and compared for target cell killing *in vitro* and *in vivo*. CAR T cells were evaluated in Incucyte® living imaging assays with co-cultures of (**A**) AsPC1 cells or (**B**) OVCAR8 cells at E:T ratios of 10:1, 3:1, 1:1 and 0.3:1 for the times indicated. Graphs were analyzed to show % killing. (**C**) Schema of CAR T cells evaluated *in vivo* in the OVCAR8 peritoneal mouse model. (**D**) CAR T cell tumor control for each of the three donors with associated mouse survival curves. ND630 CAR T cells were injected intravenously 14 days after the introduction of OvCAR8 tumor cells while ND616 and ND410 cells were injected 21 days afterwards. ND630 and ND616 CAR T groups had 7 mice/group while the ND410 groups had 10 mice/group.

We next compared these stump CAR T cells from three donors for *in vivo* efficacy of CAR 422 with CAR 15B6 and CAR M5 in our OvCAR8 tumor MSLN shedding mouse model (**Fig. 4C**). As seen in our earlier stump CAR screening, CAR 422 effectively controls tumor to baseline levels by three weeks post stump CAR T cell administration (**Fig. 4D**). The lack of efficacy in our distal targeting CAR M5 is also consistent with our earlier screening results. Our stump CAR 422 and stump CAR 15B6 T cells consistently have similar efficacy in tumor control and long-term survival.

We evaluated the engraftment level of the ND616 human T cells in the blood, spleen and peritoneal ascites fluid on day 38 after intravenous injection, when tumors had been cleared. T cells were not detected either in the spleen or blood but only in the peritoneal cavity (**Fig. S7**). This indicates that the T cells migrated into the peritoneum and remained there in the highest numbers after intravenous injection. While the CAR 422 T cell group had similar numbers in ascites as CAR M5, the M5 had lost efficacy to the MSLN, presumably due to cleavage of the targeted epitope. Interestingly, the spleen did not have residual T cells in all groups, but consistently, there was less splenomegaly in mice treated with CAR T cells targeting mesothelin (1.0 cm length) compared to the groups treated with non-targeting CAR 119 and the no treatment group (∼1.6 cm length).

### Specificity of scFv ST4-22 for MSLN

Having identified CAR 422 as our lead candidate to advance into the clinic, we wanted to confirm monospecificity of the ST4-22 binder for MSLN. ST4-22 was expressed as a bivalent scFv using a murine Fc constant region (**Fig. 5A**) and retention of binding to the appropriate form of MSLN was confirmed by ELISA (**Fig. S8**). To assess off-target binding, bivalent scFv ST4-22 was submitted to Integral Molecular, Inc. to run on their Membrane Proteome Array^TM^, a proprietary protein array comprising over 6,000 distinct human membrane protein clones each overexpressed in live cells from expression plasmids (32). The platform uses flow cytometry to directly detect antibody binding to human membrane proteins which includes 94% of all single-pass, multi-pass, and GPI-anchored proteins, GPCRs, ion channels, and transporters. Proteins are expressed in unfixed cells in their native conformation with appropriate post-translational modifications. The results of the proteome array screening with ST4-22 expressed graphically **(Fig. 5B)** indicated that ST4-22 has exquisite specificity for human MSLN (Uniprot Q13421).

**Fig. 5.**
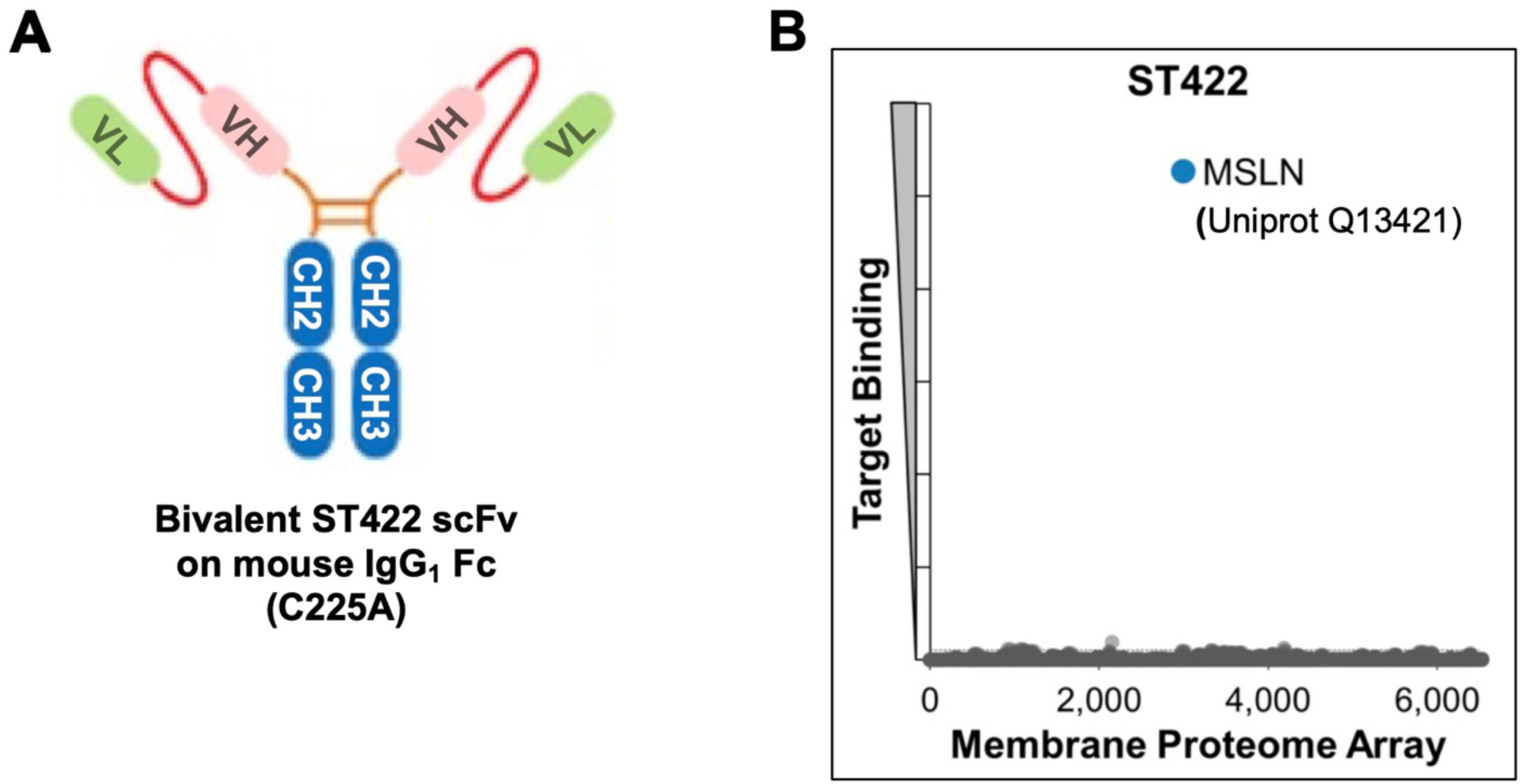
Specificity profiling of ST4-22 by testing cross-reactivity against a human membrane proteome array. (**A**) A bivalent scFv construct of ST4-22 was submitted to Integral Molecular, Inc. to probe their >6000-membrane human membrane proteome array. (**B**) Proteome screening demonstrated monospecificity for human mesothelin (Uniprot sequence Q13421).

## Discussion

CAR T cells and other approaches targeting mesothelin have promise yet face challenges of on-target off tumor toxicity and antigenic shedding (24, 29, 33–35). Our development of CAR 422 T cells targeting the juxtamembrane “stump” region of mesothelin addresses long-standing challenges in CAR T therapy for solid tumors.

Notably, shed mesothelin acts as both a decoy receptor and an immunosuppressive barrier binding CARs before they reach the tumor cell surface and reducing therapeutic efficacy (18, 19, 36). Our CAR 422 demonstrates comparable—or superior—cytotoxicity and *in vivo* tumor activity to conventional mesothelin-targeting CARs, owing to its resilience to antigen shedding.

CAR 422’s ability to bind only the membrane-adjacent stump—ignoring soluble, shed mesothelin—mirrors findings with murine-derived 15B6 and its humanized h15B6 variant (19, 20). In patient-derived xenograft (PDX) models of pancreatic cancer, h15B6 CARs produced durable remissions where SS1-based CARs targeting the amino terminus of mesothelin failed (20). Our ST4-22 binder further validates this approach, offering a canine-derived scFv that avoids cross-competition with shed antigen and enables unimpeded tumor engagement.

Native mesothelin shedding contributes to diminished antigen density on tumor cells, further reducing CAR recognition (18). By binding the un-shed stump, CAR 422 effectively bypasses this barrier. Moreover, high concentrations of shed antigen in the tumor milieu have been associated with CAR T-cell deactivation or sequestration at extraneous sites rather than the tumor (17, 19). Our construct avoids this trap, maximizing CAR availability and function at the tumor interface.

Membrane proteome array screening confirmed that our scFv ST4-22 binder shows exquisite specificity for human mesothelin, with no detectable off-target reactivity—an essential safety feature for clinical advancement. While high-affinity CARs targeting MSLN have demonstrated potential off-tumor toxicity in preclinical lung models (37), CAR 422’s focused epitope specificity and the use of a canine-human chimera may mitigate such risks. Nonetheless, only phase I clinical testing can fully evaluate the safety of CAR T cells targeting the stump epitope. A phase I clinical trial testing CAR T cells with h15B6 scFv has recently been announced (clinicaltrials.gov NCT06885697). Although CAR 422 overcomes shedding-mediated resistance, broader challenges remain in solid-tumor CAR therapy: antigen heterogeneity, tumor infiltration, immunosuppressive microenvironments, T-cell exhaustion, and persistence (36, 38). Combining CAR 422 CAR T cells with checkpoint inhibitors, tumor microenvironment modulators, or engineered cytokine support aligns with current strategies aimed at enhancing solid tumor efficacy. Future work could explore intratumoral or regional delivery routes to concentrate activity at tumor sites and limit systemic exposure, as has been tested with intrapleural administration in mesothelioma CAR trials (6, 36, 39).

The impressive activity of CAR 422 in both tumor line-derived and PDX models, especially in tumors resistant to shed-binding CARs, highlights its translational promise. Integration of this approach into early-phase clinical studies—perhaps alongside TME-modulating agents—could open new therapeutic opportunities in mesothelin-positive ovarian and pancreatic cancers. Drawing upon prior strategies with h15B6 constructs, advanced safety designs (affinity tuning, suicide genes) and combination regimens may optimize antitumor potency while maintaining safety (20).

In summary, ST4-22-expressing CAR T cells represent a significant advance in overcoming key resistance mechanisms in mesothelin-targeted immunotherapy. By precisely targeting the juxtamembrane stump, avoiding soluble antigen traps, and maintaining specificity, CAR 422 holds potential as a next-generation therapy for challenging solid tumors.

## Materials and Methods

### Protein and immune reagents for scFv isolation and analysis

Streptavidin, neutravidin, and ABTS substrate were from Thermo Fisher Scientific (#43-4301, #31000, and #37615, respectively). Biotinylated MSLN (296–600), biotinylated MSLN (296–580), HIS-tag MSLN (296–580), and biotinylated LAIR1 were from Acrobiosystems (MSN-H82E7, MSN-H82E9, MDN-H522a, and LA1-H82E3, respectively). Anti-DNP murine IgG1 isotype control mAb was from Acrobiosystems (DNP-M1). HRP-conjugated anti-M13 murine mAb was from Sinobiologicals (11973-MM05T-H). HRP anti-mouse IgG, Fcγ fragment-specific was from Jackson Immunoresearch (115-035-071). Recombinant 15B6 (19), 30D12 anti-STEAP2 (40), and bivalent ST-422 were custom produced by Biointron Biological.

### Phage display library panning

Antibody binding fragments specific to the stump region of MSLN were isolated from a canine scFv naïve/non-immune phage display library (26, 27) using a positive/negative solid phase panning strategy utilizing full-length MSLN (296–600) and MSLN (296–580) lacking the stump domain. To minimize disruption of the native conformations of MSLN proteins that can occur by direct adsorption to polystyrene substrates, MSLN proteins with a biotinylated AviTag^TM^ were used and captured to streptavidin-coated 96-well microplates.

For each round of panning, a pair of 96-well Costar 3490 ½-area microplates were coated with streptavidin, 0.5 μg per well, at 4°C overnight (24 wells of each plate for panning round 1 and 8 wells of each plate for panning rounds 2 through 4). Plates were washed in PBS, and wells were blocked with 2% non-fat dry milk in PBS (MPBS) at 37°C for 1 h. To one plate in each pair, 36 pmol of biotinylated MSLN (296–600) was added to each streptavidin-coated well. To the other plate an equivalent amount of biotinylated MSLN (296–580) was added. After a 1-h incubation at 37°C, plates were washed with PBS to remove free MSLN proteins. Equal aliquots of μκ, μα, γκ, and γα scFv phage display libraries were combined, blocked for 1h at RT in 2% MPBS, and 50 μl added to each well of the MSLN (296–580)-coated plate. After a 1-h incubation at 37°C, negatively absorbed phage was applied to wells coated with MSLN (296–600) for positive selection on full-length MSLN. For panning rounds 2 through 4, negatively absorbed phage was additionally spiked with 1 nmol of non-biotinylated MSLN (296–580) before being added to the positive selection plate. Panning rounds and overnight phage amplification were continued as previously described (41, 42) where, after a 2-h incubation at 37°C, unbound phage were washed away 5 times during the first panning and 10 times during each subsequent panning round using PBS supplemented with 0.1% Tween 20 (PBST). Each wash was performed with a 5-min incubation of wash buffer in wells to select for binders with longer off-rates.

### ScFv-phage ELISA

Monoclonal phage-displayed scFvs were prepared from panning eluates as described (43), added to microplates previously coated with streptavidin and biotinylated proteins and blocked with 2% MPBS as in panning procedure. From experience, a 1:100 dilution of overnight monoclonal scFv-phage-containing cultures provides consistent ELISA signals suitable for positive/negative screening. After washing away unbound phage with PBST, HRP-conjugated anti-M13 mAb (1:10,000) was added for 1h at 37°C and the microplate was washed, developed with ABTS substrate solution, and read at wavelengths of 410 nm and 490 nm.

### 15B6 competition ELISA

Microplate wells were coated with 0.5 μg/well streptavidin, blocked with 2% MPBS, and loaded with biotinylated MSLN (296–600). To increase the sensitivity of the assay, one-half of the amount of MSLN protein used in panning procedure was added per well (i.e., 18 pmol). For each scFv-phage to be tested, one well of a pair of MSLN-coated wells was left filled with blocking buffer and the other was filled with 15B6 diluted in blocking buffer. A titration of 15B6 (**Fig. S2**) had indicated that under these conditions, 15B6 binding saturated at ∼1 μg/ml. Therefore, a 10-fold excess of 15B6 (10 μg/ml) was used here. Microplates were incubated for 30 min at 37°C to allow 15B6 to bind and then 1/9^th^ volume of 10x-strength scFv-phage was added to each well of a pair to bring the final scFv-phage concentration to standard ELISA phage dilution of 1:100. After an additional 1h at 37°C, wells were washed with PBST, developed with HRP-conjugated anti-M13, and developed with ABTS substrate as above.

### Nucleotide sequencing of scFv clones and analysis

Mini-scale DNA plasmid preparations of the pComb3X phagemid vector were made from monoclonal scFv-phage cultures, and nucleotide sequences for the heavy and light chain scFv variable regions were determined using the pComb3X vector-specific sequencing primers “5’LC” and “dpseq” that anneal just upstream of the light chain and downstream of the heavy chain variable region sequences, respectively (25).

Sequencing was performed by the University of Pennsylvania Department of Genetics DNA Sequencing Core Facility using the di-deoxynucleotide chain termination (Sanger) method on an ABI 3730xl Sequencer.

Nucleotide sequences were analyzed using MacVector software (v. 18.7.8). The online IMGT/V-QUEST germline gene analysis tool (44) available at http://www.imgt.org/IMGT_vquest/input and the online IMGT/DomainGapAlign immunoglobulin protein analysis tool (45) available at http://www.imgt.org/3Dstructure-DB/cgi/DomainGapAlign.cgi were used together to identify V, D, and J germline genes and identify framework regions and CDRs.

### Membrane proteome array screening

Bivalent scFv ST4-22 was analyzed on the Integral Molecular Membrane Proteome Array^TM^ (32). Each of over 6,000 distinct membrane protein clones was individually transfected in separate wells of 384-well plates followed by a 24-h incubation. Cells expressing each individual protein clone were arrayed in duplicate in a matrix format for high-throughput screening. Before screening on the array, the ST-422 concentration for screening was determined on cells expressing positive (membrane-tethered Protein A) and negative (mock-transfected) binding controls, followed by detection by flow cytometry using a fluorescently labeled secondary antibody. ST-422 was added to the array at the predetermined concentration, and binding across the protein library was measured on an Intellicyt iQue using an AlexaFluorR 647 labeled goat F(ab’)2 anti-mouse Fc secondary antibody (Jackson ImmunoResearch 115-606-008). Each array plate contained both positive (Fc-binding) and negative (empty vector) controls to ensure plate-by-plate reproducibility. Test ligand interactions with any targets identified by membrane protein array screening were confirmed in a second flow cytometry experiment using serial dilutions of the test antibody, and the target identity was re-verified by sequencing.

### Incucyte® assays

eGFP transduced tumor cells (AsPC1, H838, OVCAR8) were cocultured with CAR^+^ T-cells at various effector to target ratios. CAR killing activity was quantified by decrease in GFP fluorescent intensity using Incuyte® SX5’s Adherent Cell-by-Cell analysis software. Incucyte® Cytotox NIR Dye was added at 1:200 dilution to each well to monitor cell death. Signal intensities were quantified every three hours for seven days.

### Chronic antigen exposure (CAE) stress test

GFP positive AsPC1 cells were cocultured with CAR T-cells at a 1:1 effector to target ratio. Incucyte® Cytotox Dye for Counting Dead Cells was added at 1:200 dilution to each well. CAR killing activity was monitored by the Incucyte® every 3 hours. CAR T-cells were restimulated with new AsPC1 cells when 50% killing was achieved. At restimulation, the media and T-cells were collected and spun down. Fresh AsPC1 cells were re-seeded onto the plate. Half of the conditioned media and all T-cells were added back to the well. Restimulation was repeated until the exhaustion profile was characterized.

### CAE flow analysis

At each restimulation, cells were removed for flow analysis of CAR T-cell number and phenotype. Each CAR T-cell was evaluated for viability with LIVE/DEAD™ Fixable Aqua Dead Cell Stain Kit (Thermo Fisher Scientific, # L34966) then stained for surface CAR expression with Biotin-SP AffiniPure® F(ab’)₂ Fragment Goat Anti-Human IgG (Jackson ImmunoResearch, #109-066-006), Biotin-SP AffiniPure® Goat Anti-Mouse IgG (Jackson ImmunoResearch, #115-065-072), or Rabbit Anti-Dog IgG F(ab’)₂ Biotin Conjugated Affinity purified (Rockland, #604-4604), followed by PE-Streptavidin (BioLegend, #405203) secondary staining. The following human markers were measured: CD45 (BioLegend, #304011), CD4 (BioLegend, #317442), CD8 (BioLegend, #3009100), PD-1 (BioLegend, #329928), Tim3 (BioLegend, #345018), TIGIT (BioLegend, #372714), KLRG1 (BioLegend,#368604). CountBright Absolute Counting Beads (Invitrogen, #C36950) were added to each sample. Samples were run on a BD LSR Fortessa. Data was analyzed with FlowJo v10.10.0.

### Tumor cell lines

AsPC1, a pancreatic adenocarcinoma cell line (ATCC CRL-1682), H838, a non-small cell lung adenocarcinoma (ATCC CRL-5844), and OVCAR-8, an ovarian adenocarcinoma (RRID:CVCL-1629) were each transduced to express eGFP-2A-CBR for monitoring MSLN CAR targeting. MSLN expression was knocked out in AsPC1 and H838 cells using CRISPR technology for evaluating non-MSLN activity of the stump CARs. These MSLN KO cells were transduced to express only the stump region with a 5’ FLAG-tag via GPI linkage for confirming stump CAR regional binding specificity.

### Engineering human CAR T cells

Lentiviral constructs encoding CAR transgenes were packaged, titered, and transduced into human T-cells activated by CD3/CD28-Dynabeads (Thermo Fisher Scientific, #43500D). Transduced T-cells were cultured and expanded with X-VIVO 15 (Lonza Biosciences, #02-060Q) and 5% HuAB serum supplemented with IL7 and IL15 at 5 ng/mL. CAR transduction efficiency was determined by flow cytometry and the cells were expanded for 10 days before freezing.

### Mouse models

For the AsPC1 subcutaneous model, 2e6 AsPC1 cells were suspended in 50% Matrigel (Corning, #356234) and injected subcutaneously into NOD scid gamma (NSG) mice.

Mice were treated with CAR T cells when tumors reached ∼200 mm^3^ in volume after about two weeks. Tumor sizes were measured weekly.

For the OVCAR8 peritoneal model, 3e6 OVCAR8 cells were injected into the peritoneal cavity of NSG mice and allowed to develop for about 3 weeks before CAR T-cell IV injection. Tumor progression was measured weekly by bioluminescence using the IVIS Lumina system and data was analyzed using Living Image®.

Pre-specified endpoints of animal experiments were the health of animals and tumor burden. All procedures followed the guidelines of an IACUC approved animal protocol at the University of Pennsylvania.

### Ascites collection and analysis

Ascites fluid was collected by injecting 3% FBS buffer into the peritoneal cavity, incubated for 10 minutes, and collected using 5-mL syringes. The collected ascites was analyzed for CAR T-cell persistence and their phenotypic profile and soluble mesothelin levels.

### Ascites flow staining

Cells in ascites had red blood cells removed with ACK RBC lysis buffer (Lonza Biosciences, #BP10-548E). The resulting cells were stained for the following human markers CD45 (BioLegend, #368516), CD4 (BioLegend, #317434 or #317434), CD8 (BioLegend, #344742), CD45RO (BioLegend, #304210), CD62L (BioLegend, #304822), HLA-DR (BioLegend, #327022), Tim3 (BioLegend, #345006), PD-1 (BioLegend, #329928) and CD39 (BioLegend,#328224). Mouse myeloid markers evaluated were CD45 (BioLegend, #368516) and CD11b (BioLegend, #550993).

## Acknowledgments

We would like to thank Khatuna Gabunia, Mei “Vivian” Ji, and Ting-Jia Fan for technical assistance and/or helpful discussions and Krishna Vijayendran for contributions to Figure 1. Figure 1 is a modification of that presented in Ma et al. (46) and is subject to the terms of the Creative Commons Attribution License at creativecommons.org/ licenses/by/4.0/. We acknowledge the Apheresis Unit and Human Immunology Core (HIC) at the University of Pennsylvania for a reliable supply of healthy human T cells.

Funding was provided by 1R01CA287866 and P01 CA214278 (C.H.J.).

---

**Table S1.**
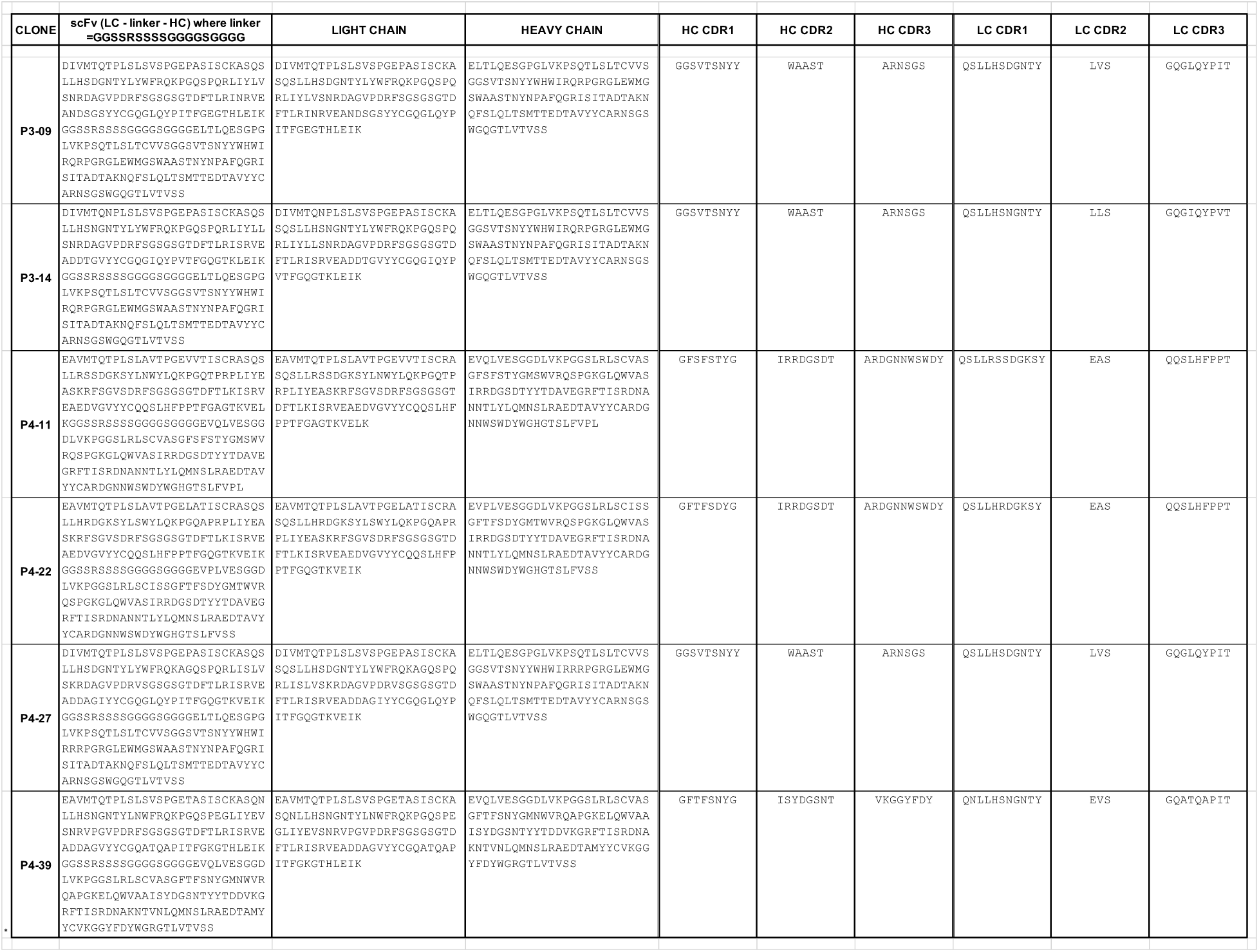
Amino acid sequences of MSLN stump-binding scFvs.

**Table S2.**
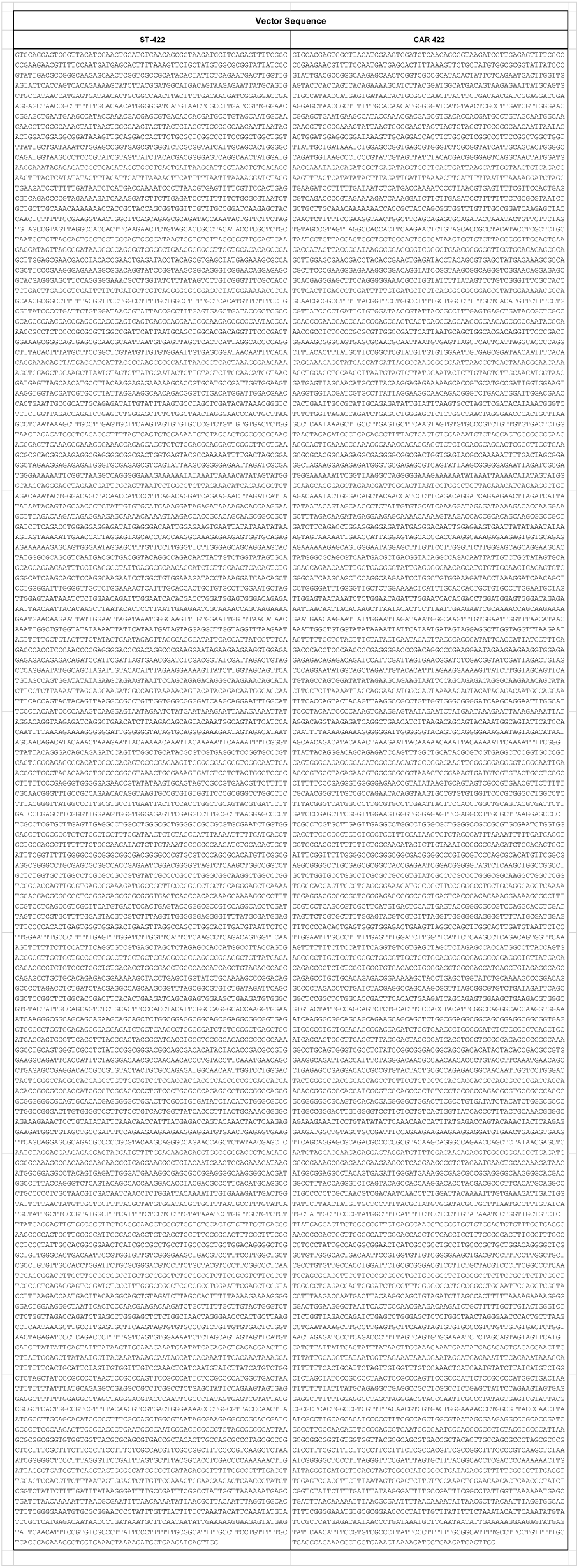
Nucleotide sequences for vector before (ST-422) and after (CAR 422) ORF optimization.

**Fig. S1.**
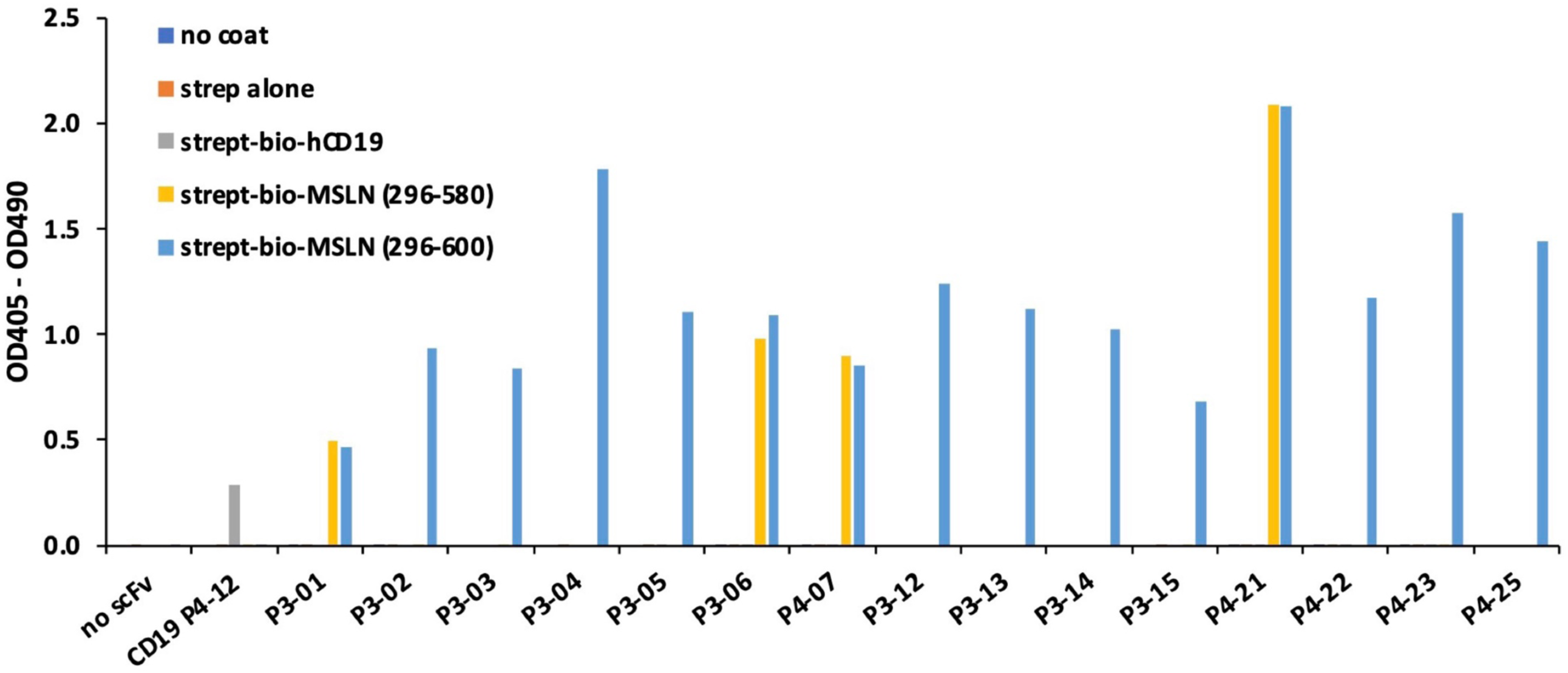
MSLN binding patterns of scFvs derived from phage library panning campaign. Representative phage ELISA showing binding of 15 (of 66) randomly selected scFv clones to MSLN (296–600) and MSLN (296–580) polypeptides representing full-length and shed domains, respectively. Results for this representative group show that P3-01, P3-06, P4-07, and P4-21 bind both MSLN polypeptides suggesting specificity for shed MSLN while the rest bind only MSLN (296–600) which contains the stump domain. Controls included uncoated microplate wells and wells coated with only streptavidin (strep) or streptavidin and biotinylated human CD19 protein, as well as a phage displayed anti-human CD19 scFv isolated from the canine phage library in a different panning campaign.

**Fig. S2.**
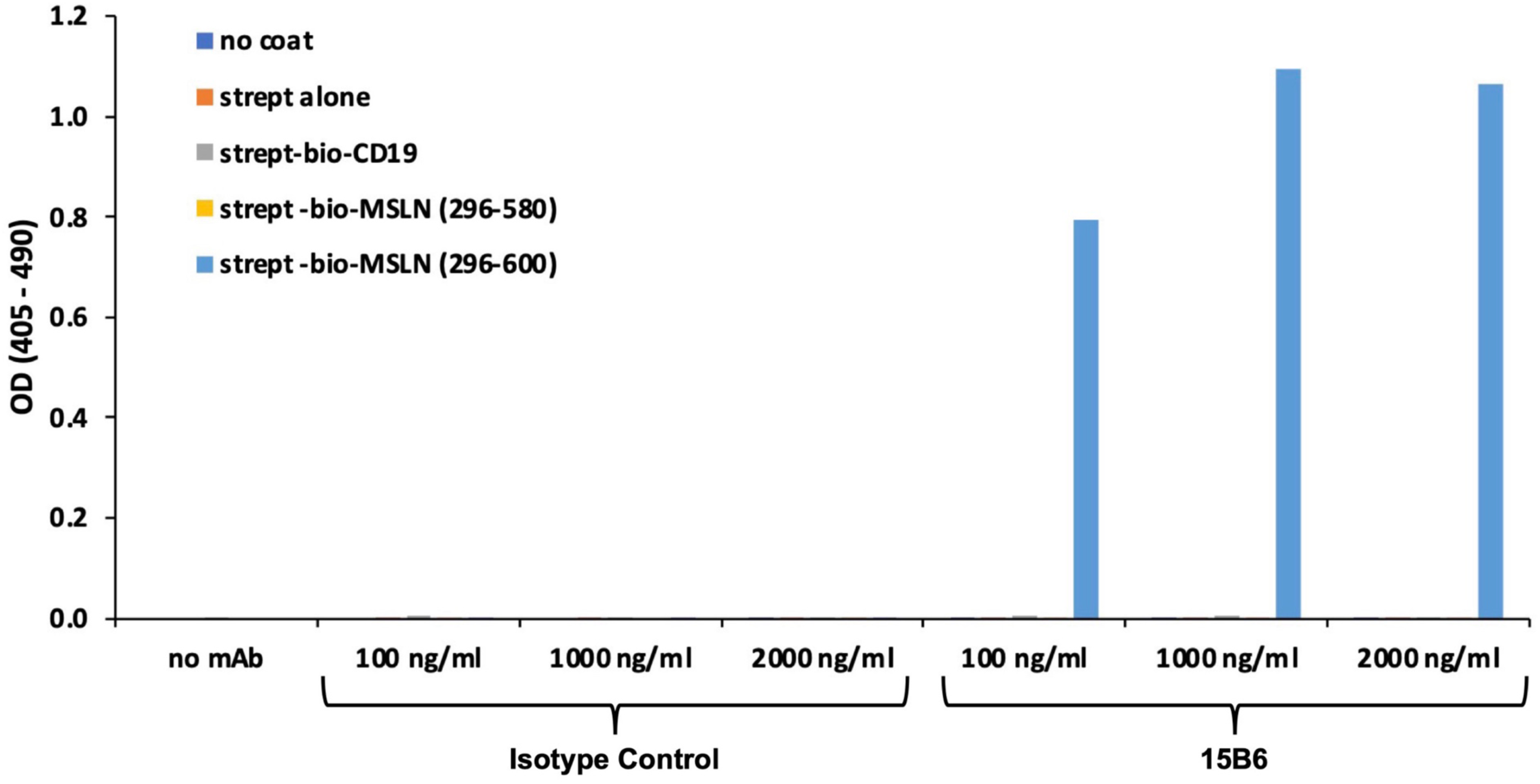
Binding of recombinant murine 15B6 to MSLN polypeptides. ELISA to verify that recombinantly produced 15B6 reference antibody retains binding to MSLN stump region. Microplate wells were uncoated or coated with streptavidin (strept) alone, or streptavidin and biotinylated human CD19, MSLN (296–580), and MSLN (296–600) and reacted with recombinant 15B6 (or a murine isotype control (anti-dinitrophenol)). Recombinant 15B6 binds only stump-containing full-length MSLN (296–600) but not truncated MSLN (296–580).

**Fig. S3.**
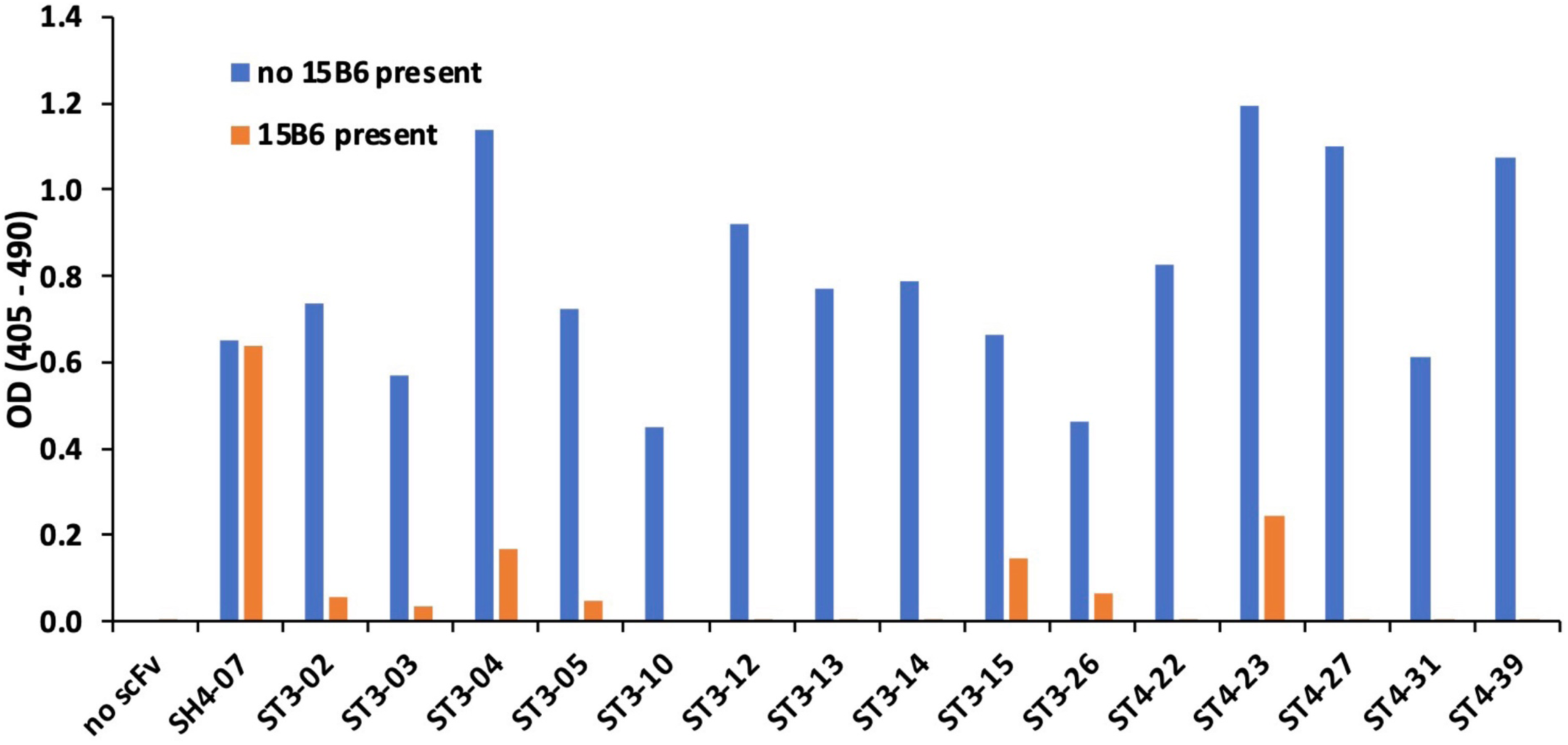
15B6 competition ELISA. Representative MSLN (296–600) binding profiles shown for 15 of 29 stump-binding phage display-scFvs in the presence or absence of pre-bound 15B6 reference antibody. Ratios of binding in the absence to the binding in the presence of 15B6 were calculated from these data for all 29 scFvs and listed in **Fig. 2C**. Internal control scFv SH4-07, shown to be directed to shed MSLN polypeptide (**Fig. S1**), shows no inhibition by stump-directed 15B6 as expected.

**Fig. S4.**
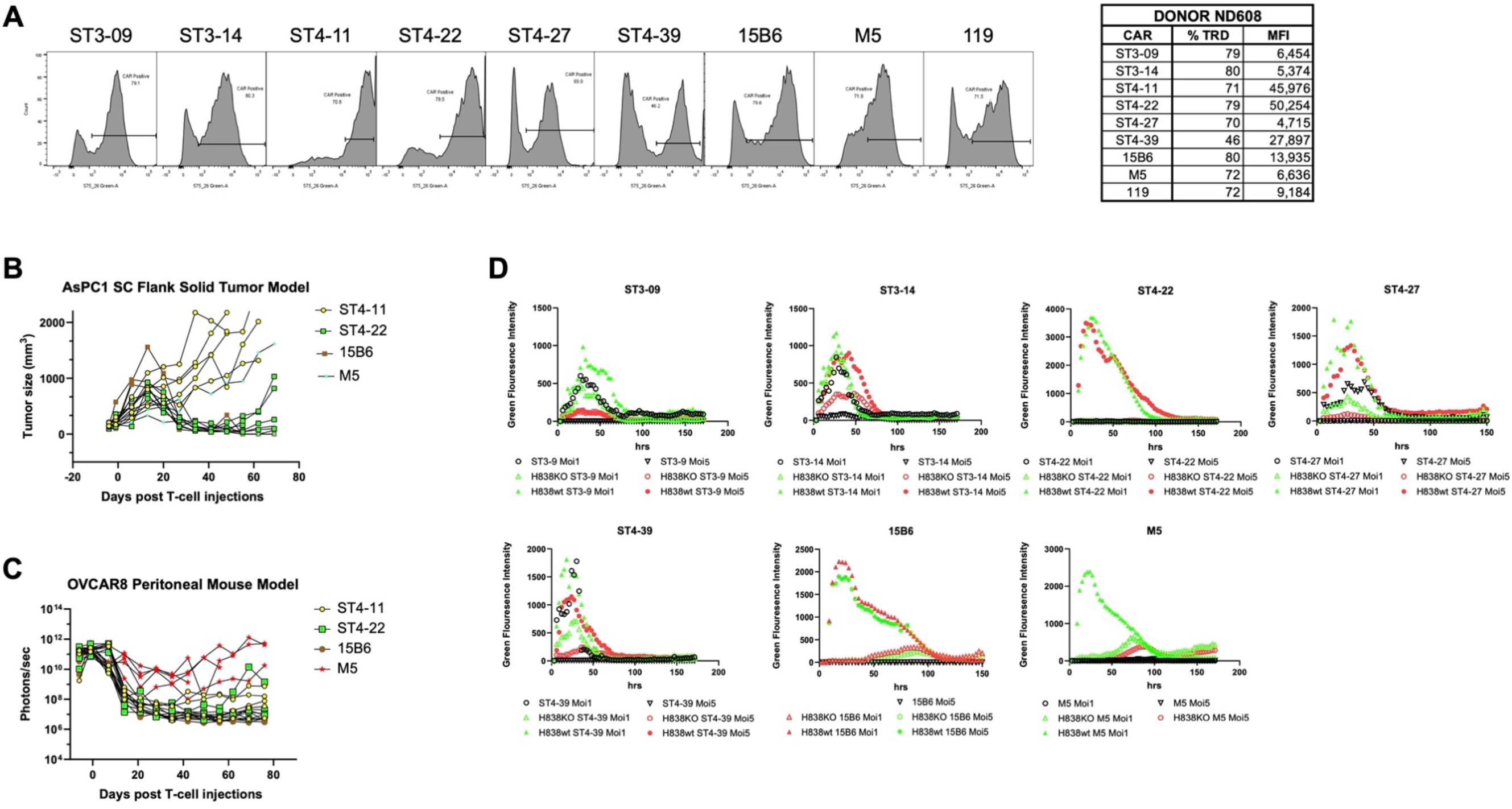
Expression and activation properties of MSLN stump-directed CAR-T cells. (**A**) Flow histograms of healthy donor ND608 T cells showing expression of MSLN-directed and CD19-directed (119, control) CARs. Percent transduction (% TRD) and mean fluorescence intensity (MFI) are tabulated. (**B**) AsPC1 tumor control profiles of individual mice treated with selected CAR T cells to illustrate scatter of responses. (**C**) OVCAR8 tumor control profiles of individual mice treated with selected CAR T cells to illustrate scatter of responses. (**D**) Peak activation assay showing basal (tonic activity) and activation kinetics of Jurkat-NFAT GFP reporter cells transduced with MSLN-directed CAR T cells and co-cultured alone or with H838 wild type or MSLN-KO cells. Co-cultures were either at a 1:1 or 5:1 E:T ratio.

**Fig. S5.**
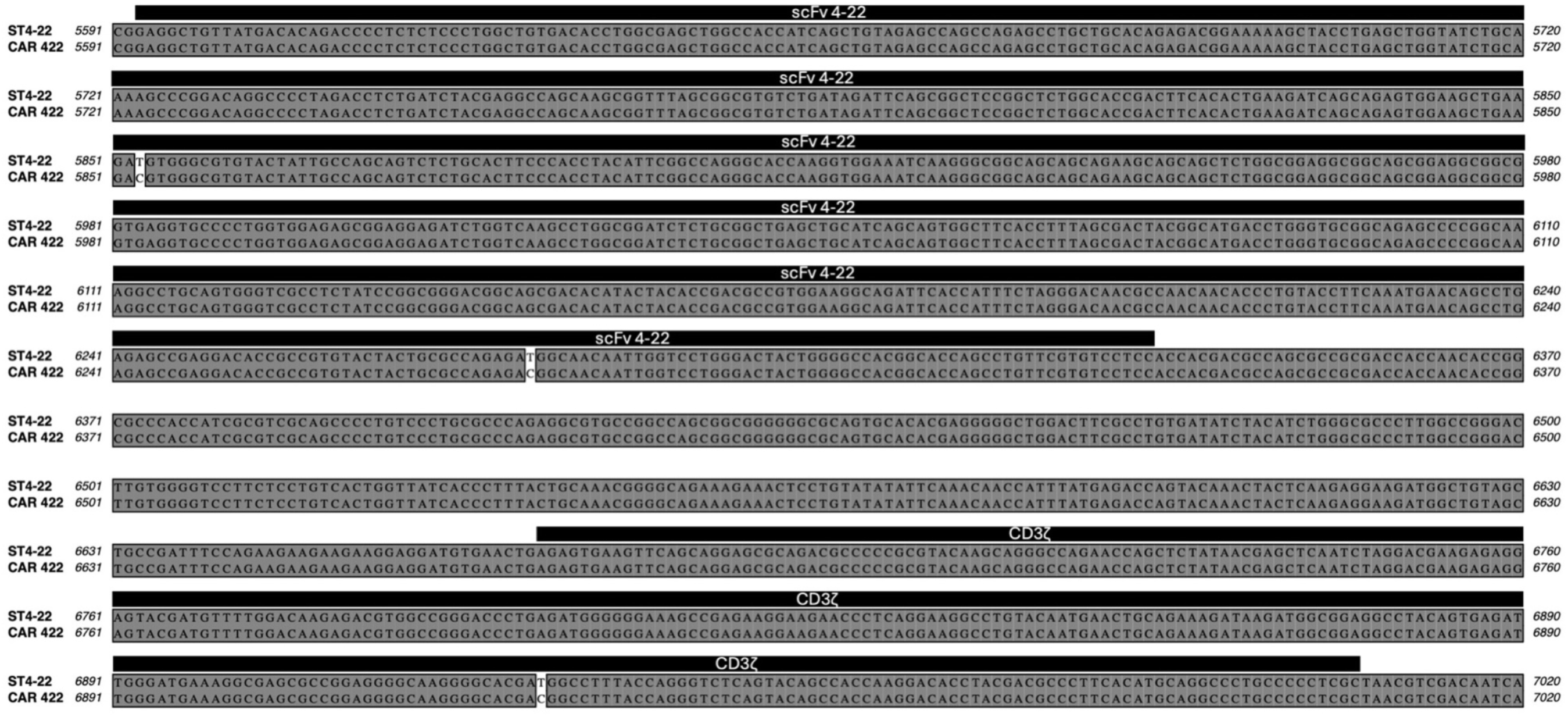
Nucleotide alignment of CAR vectors for ST4-22 and CAR 422 to show elimination of internal ATG start sites from alternate reading frames. Locations for the 3 modifications within vector nucleotide positions 5591 and 7020 are shown along with positions encoding scFv 4-22 and CD3ζ domain.

**Fig. S6.**
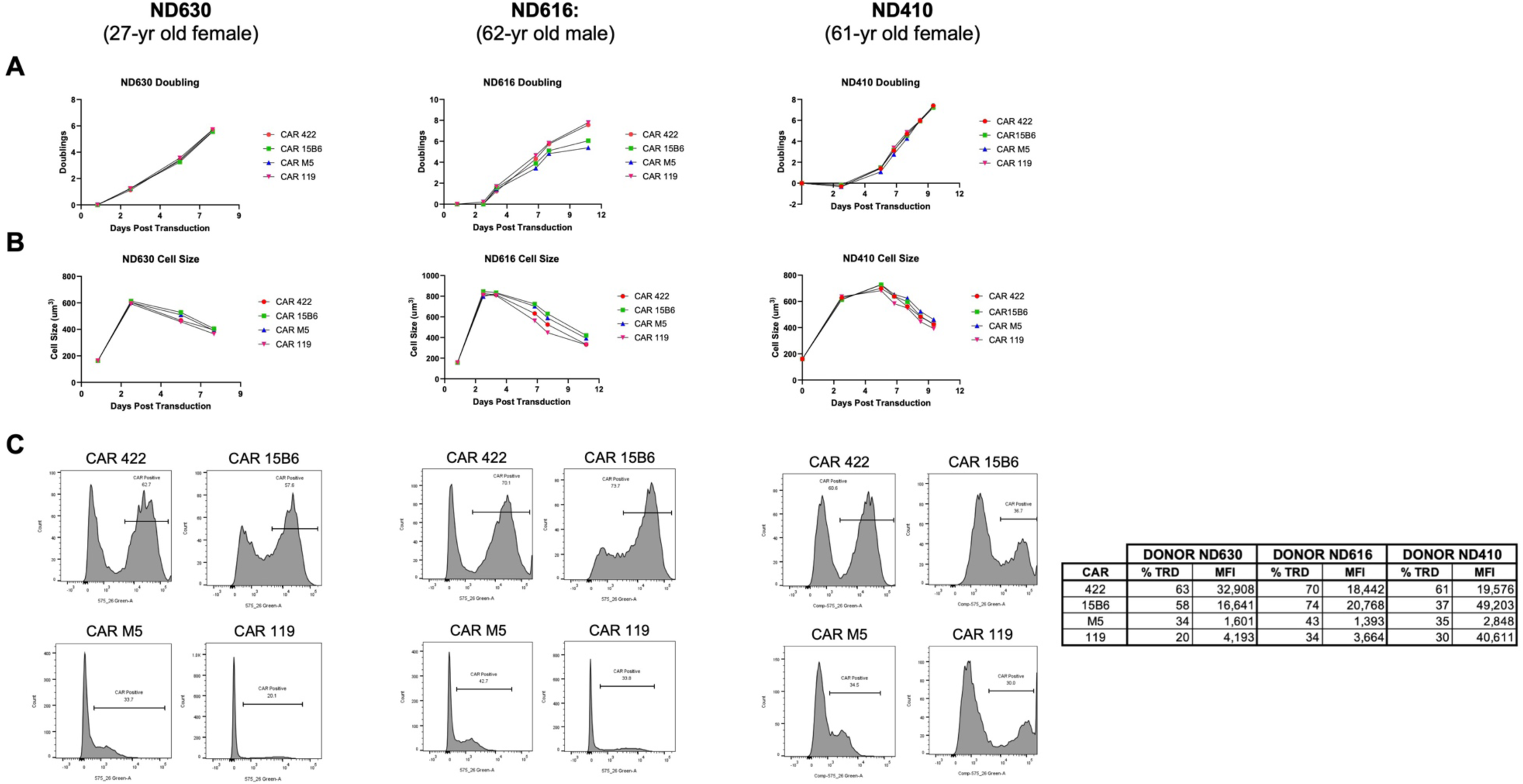
CAR T cell manufacturing profiles in cells from three healthy human donors. MSLN-directed CARs 422, 15B6, and M5, and CD19-directed CAR 119 were transduced into three healthy donors ND630, ND410 and ND616 and their (**A**) rate of doubling and (**B**) cell size are compared. (**C**) Percent transduction efficiency (% TRD) and mean fluorescent intensity (MFI) are displayed in histograms and tabulated.

**Fig. S7.**
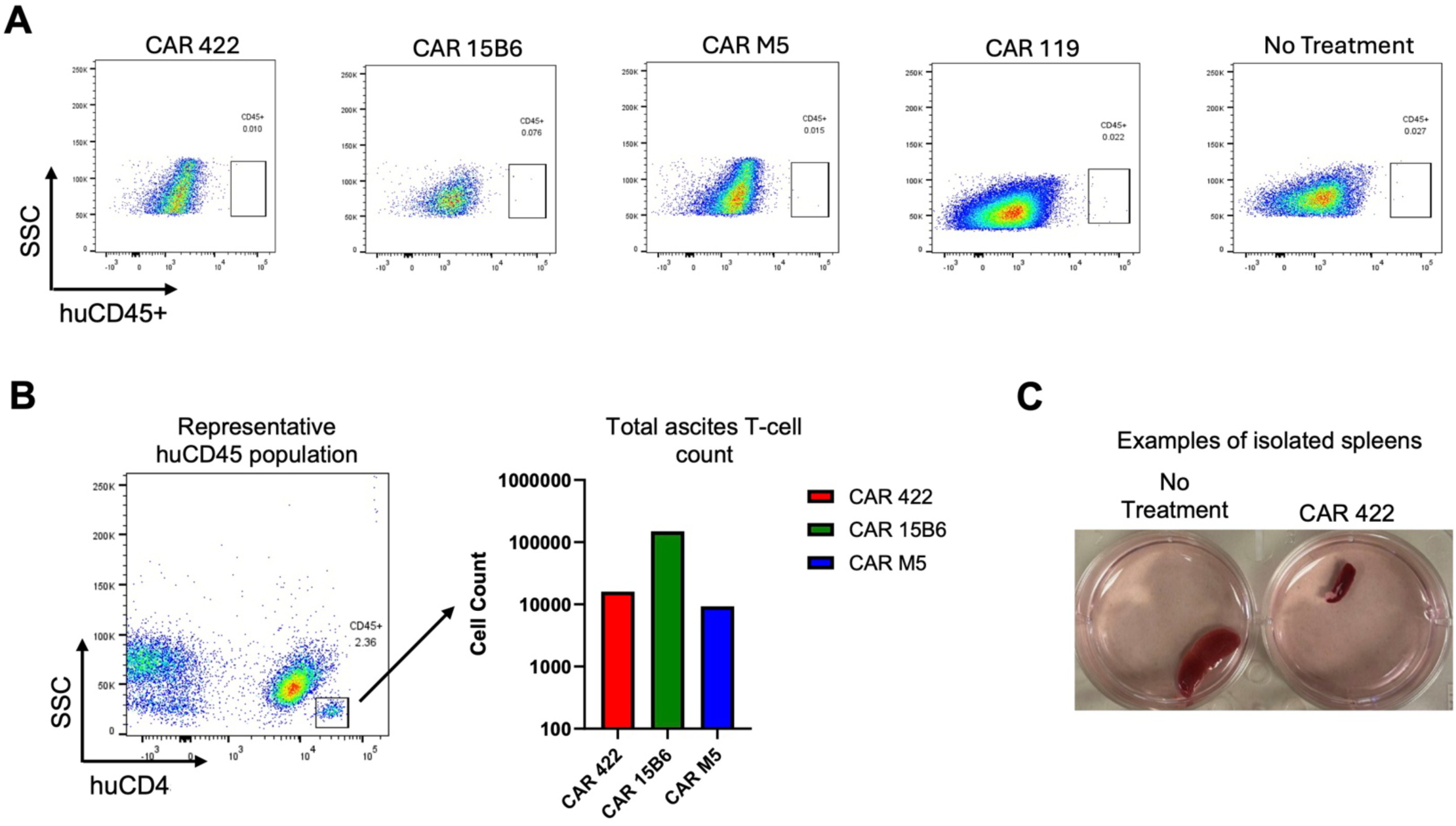
Engraftment levels of CAR T cells in the OVCAR8 peritoneal mouse model. After tumors had cleared by day 38 of the OVCAR8 experiment (**Fig. 4C, D**), mice from IV-administered ND616 CAR T cells (3 mice from each CAR-T group) were sacrificed, and their blood, spleen, and ascites were evaluated for the presence of human T cells by flow cytometry. (**A**) Examination of peripheral blood and spleen reveals absence of human T cells. (**B**) Examination of ascites reveals migration of human T cells into the peritoneum. (**C**) Representative images of mouse spleens showing a consistently smaller size in MSLN-targeting CARs than in control groups.

**Fig. S8.**
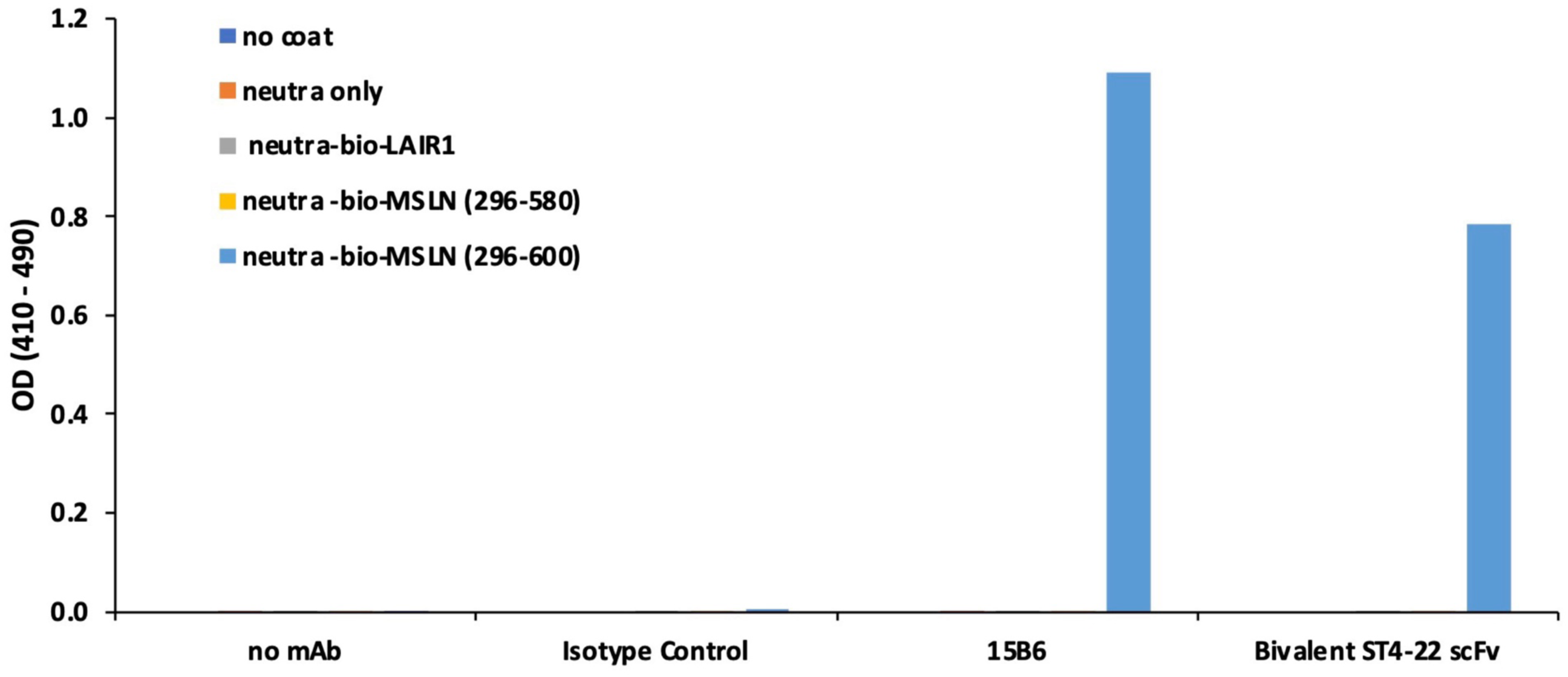
Binding of bivalent ST422 scFv to MSLN polypeptides. A bivalent scFv form of ST4-22 produced to assess off-target binding in a membrane proteome array was first validated for its binding to MSLN (296–600) and MSLN (296–580) polypeptides representing full-length and shed domains, respectively. Controls included uncoated microplate wells and wells coated with only neutravidin (neutra) or neutravidin and biotinylated human LAIR1 protein. Bivalent ST4-22 scFv along with an isotype control (murine anti-STEAP2) and reference antibody 15B6 were applied to wells at 2000 ng/ml. Bivalent ST4-22 scFv binds only to full-length MSLN but not to polypeptides representing shed MSLN as expected.

